# A comprehensive computational investigation into the conserved virulent proteins of *Shigella* sp unveils potential siRNA candidates as a new therapeutic strategy against shigellosis

**DOI:** 10.1101/2021.03.09.434519

**Authors:** Parag Palit, Farhana Tasnim Chowdhury, Namrata Baruah, Bonoshree Sarkar, Sadia Noor Mou, Mehnaz Kamal, Towfida Jahan Siddiqua, Zannatun Noor, Tahmeed Ahmed

**Author notes:** Corresponding author: Zannatun Noor, PhD.

## Abstract

*Shigella* sp account for the second-leading cause of deaths due to diarrheal diseases among children of less than 5 years of age. Emergence of multi-drug resistant *Shigella* isolates and the lack of availability of *Shigella* vaccines have made the efforts in the development of new therapeutic strategies against shigellosis very pertinent. In our study we have analyzed a total of 241 conserved sequences from a 15 different conserved virulence genes of *Shigella* sp and through extensive rational validation using a plethora of computational algorithms; we primarily obtained fifty eight small-interfering RNA (siRNA) candidates. Further extensive computational validation showed only three siRNA candidates that were found to exhibit substantial functional efficacy, be non-immunogenic and have a thermodynamically stable and sterically feasible and thereby acceptable tertiary structure. These siRNA candidates are intended to suppress the expression of the virulence genes, namely: IpgD (siRNA 9) and OspB (siRNA 15 and siRNA 17) and thus act as a prospective tool in the RNA interference (RNAi) pathway. However, the findings of our study require further wet lab validation and optimization for regular therapeutic use in the treatment of shigellosis.

## Introduction

Shigellosis can be attributed for approximately 12.5% cases of diarrheal mortality on a global scale; accounting for about 163400 cases of annual deaths, 54900 of these cases being children of less than 5 years of age (1). Global burden of shigellosis is mainly attributable to *S. flexneri* and *S. sonnei* whereas, *S. boydii* is uncommon outside South-east Asia (2). In regions with rapid industrialization, development in water sanitation and in economically developing regions, *S. sonnei* has gained prevalence in the epidemiological shift in places like Vietnam, Thailand and Bangladesh (3–6). *S. sonnei* is also predominant in traveler’s diarrhea (7).

Although antibiotic resistant strains are being reported against ciprofloxacin, azithromycin, ceftriaxone are continued to be utilized as mainstay treatment strategies (8). The growing rise of antimicrobial resistance to *Shigella* sp. has thus led to the need for development of newer therapeutic strategies against Shigellosis more pertinent (9–15). A multi-centric study reported that 85% of cases occurring in LMIC could be attributable to *S. flexneri* 2a, 3a and 6 together with *S. sonnei* and therefore, a quadrivalent vaccine targeting these strains is expected to provide significant protection in endemic regions (16). Serotype based vaccines include conjugate vaccine, carbohydrate vaccine, live-attenuated or killed whole cell vaccines (17–20). However, due to the lack of ideal animal models, no *Shigella* vaccine has reached the stage of commercialization till date (21).

The concept of designing a single therapeutic candidate with substantial efficacy and potency against all *Shigella* sp would thereby involve targeting the multiple conserved proteins involved in the process of invasion and pathogenesis. The virulence-related proteins or Vir proteins such as VirA helps in entry and intracellular motility, golgi fragmentation in host (22, 23), VirB activates the invasion proteins (24), VirF, involved in cell invasion and activation of VirG and VirB (25). VirG, also called IcsA, helps in actin polymerization and hence movement of the bacteria from one cell to another (26). The invasion plasmid antigens, Ipa, are the key regulators of invasion and pathogenicity and also activate other related proteins (27–30). IpgB is an invasion effector protein of the Type III Secretion system (T3SS), involved in membrane ruffling for cellular entry of *Shigella* (31). IpgD is another effector involved in entry and survival of the bacteria (32–34). There are many membrane excretion proteins and surface presentation antigens together called as Mxi-Spa proteins that make up the T3SS and help in pathogenesis and invasion. In particular, Spa 33 belonging to this class of gatekeeper proteins controlling T3SS protein secretion into host cells, helps in the secretion of Ipa proteins (35), while MxiC which interacts with the Ipa proteins to mediate the entry and survival of the bacteria (36). Other T3SS effectors such as OspB, OspF and OspG modulate host cell signaling pathways, transcription and manipulate the host inflammatory response (34, 37, 38).

Recent developments in the quest for newer therapeutic strategies against a number of infectious pathogens involves the utilization of the concept of post-transcriptional gene silencing by designing siRNA molecules specific to the pathogen (39). Small-interfering RNA (siRNA) represent a class of exogenous double-stranded non-coding RNA that binds with specific sequences of a messenger RNA (mRNA) and promotes the process of degradation of the mRNA, eventually halting the process of transcription (40). Concurrent works involving the use of a plethora of computational tools and algorithms have been design potential therapeutic siRNA candidates against a number of infectious organisms, including *P.vivax* (41) and *L.donovani* (42), the COVID-19 pandemic causing SARS-CoV-2 (43) and against a number of bacterial pathogens, such as: *M. tuberculosis* (44), *Salmonella* (45) and *Listeria* (46).

In this study, we have aimed to design novel therapeutic options against all members of the *Shigella* genus that are pathogenic to humans (*S.flexneri, S.dysenteriae, S.sonnei* and *S.boydii*) through the design of siRNA candidates against a number of conserved *Shigella* proteins, involved in the process of invasion and pathogenesis. We have targeted a total of 15 conserved virulent proteins of *Shigella* sp, including: IcsA/Vir G, IpaA, IpaB, IpaC, IpaJ, IpgB, IpgD, MxiC, OspB, OspF, OspG, Spa33, VirA, VirB and Vir F and through the use of rigorous computational tools and algorithms to select the most feasible and effective siRNA candidates against all the *Shigella* sp.

## Results

### Retrieval of nucleotide sequences of conserved virulent proteins of *Shigella* sp

A total of 241 conserved sequences (shown in **Supplementary File 1**) from the 15 different conserved virulent *Shigella* proteins were obtained from subsequent multiple sequence alignment analysis, done using Clustal Omega.

### Designing siRNA candidates by using a combination of first-generation and second-generation algorithms

siDirect 2.0, a highly efficient web based computational tool was used to predict potential target specific siRNA candidates on the basis of three first generation algorithms governing the sequence preference of siRNA, namely: Ui-Tei, Amarzguioui and Reynolds rules (U, R, A rules) (47). This tool predicted a total of 58 potential siRNA candidates from the 241 conserved regions of the 15 conserved virulent *Shigella* proteins, which were found to comply with the three first generation algorithms for siRNA designing, i.e.-U, R, A rules.

Out of these 58 siRNA candidates that fulfilled the U, R, A rules, only 38 siRNA candidates were found to comply with the 90% threshold set by the i-SCORE Designer software for the second generation algorithms for siRNA designing (48). Table 1 illustrates the results of the analysis obtained from the first and second generation algorithms of siRNA designing. NCBI BLAST program was used to confirm off-target resemblance of siRNAs and none of the 38 siRNA candidates that had fulfilled both the first and second-generation algorithms for siRNA designing had displayed any off-target similarity with the human genome.

**Table 01:**
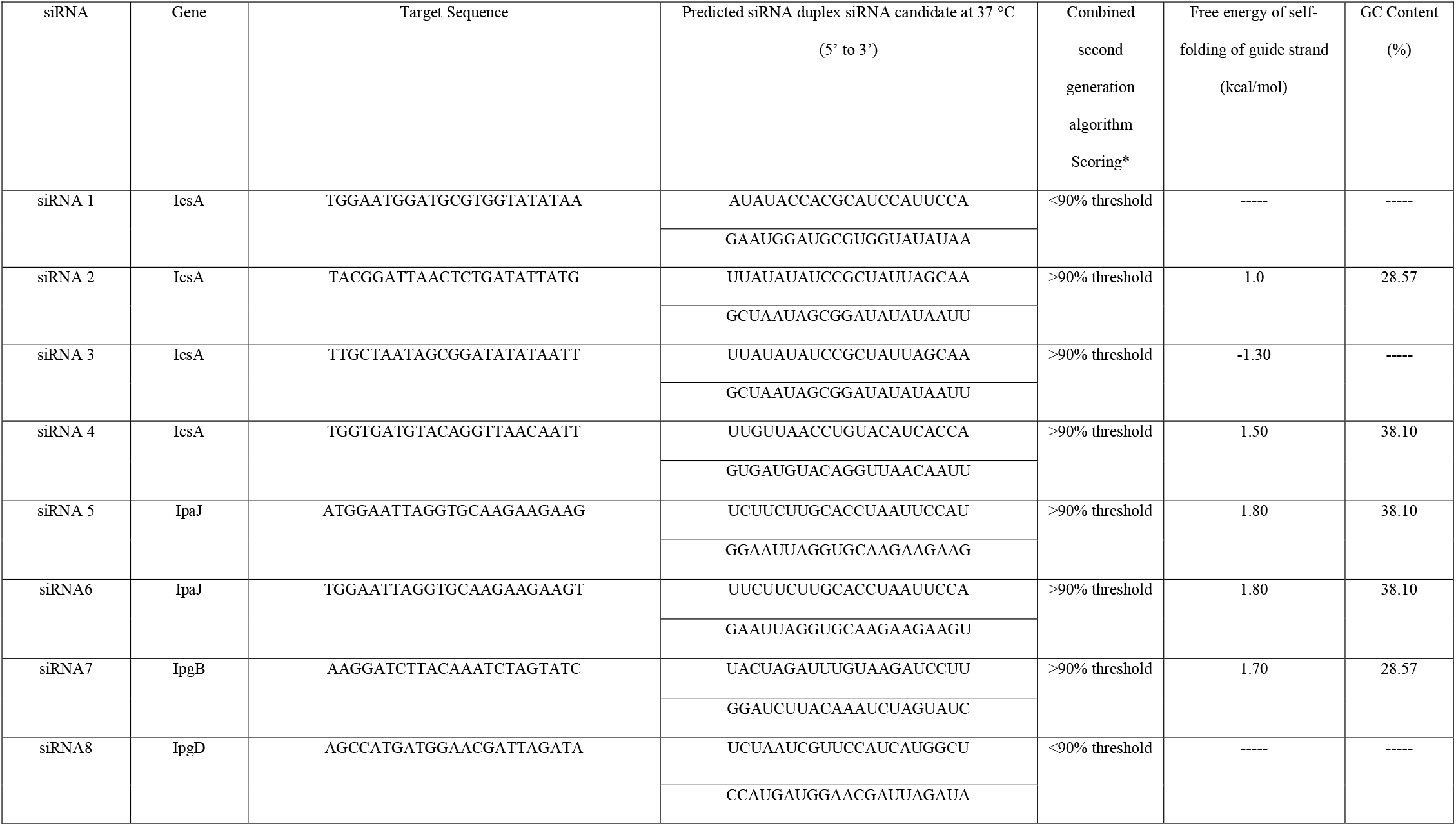
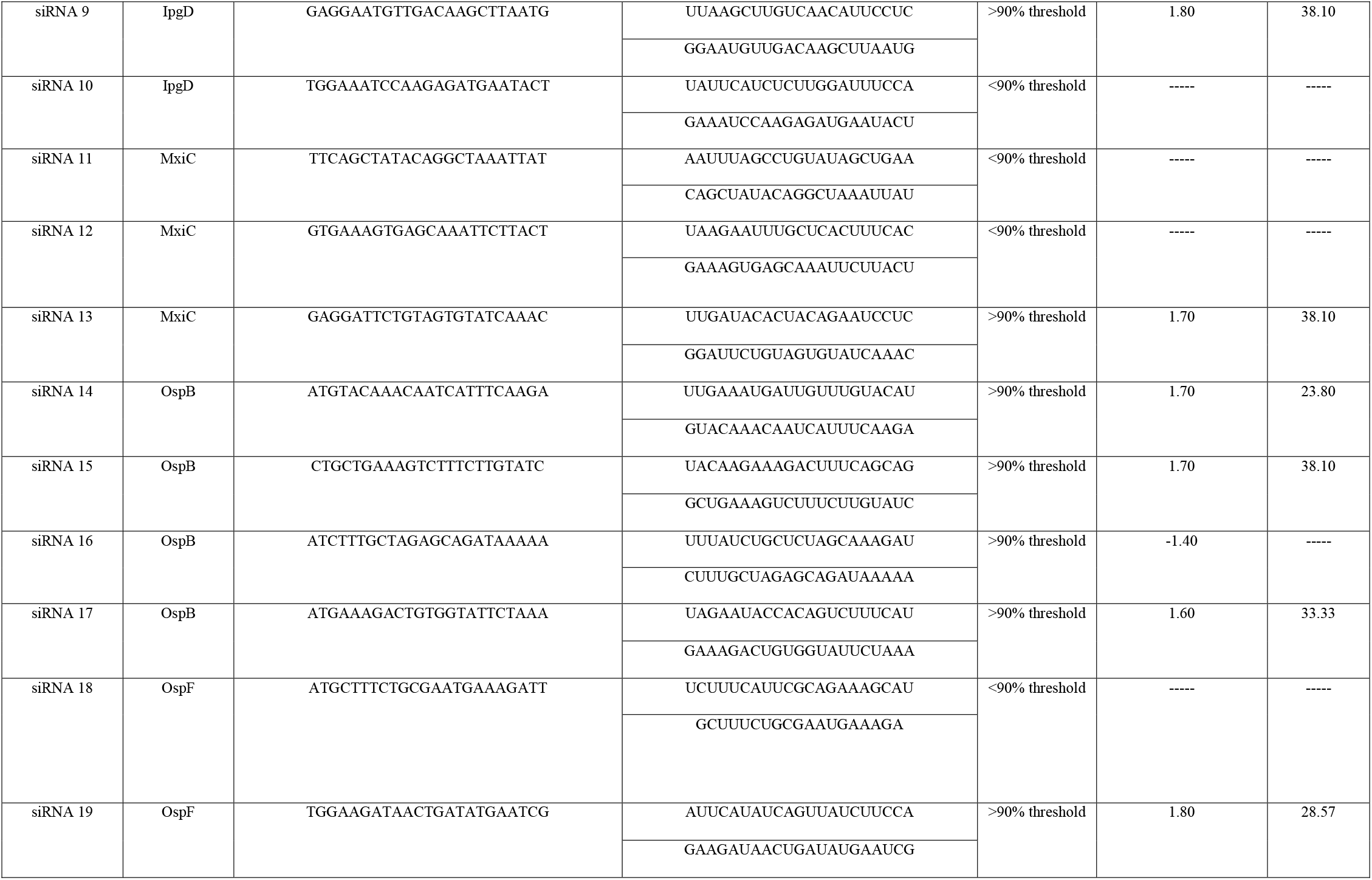
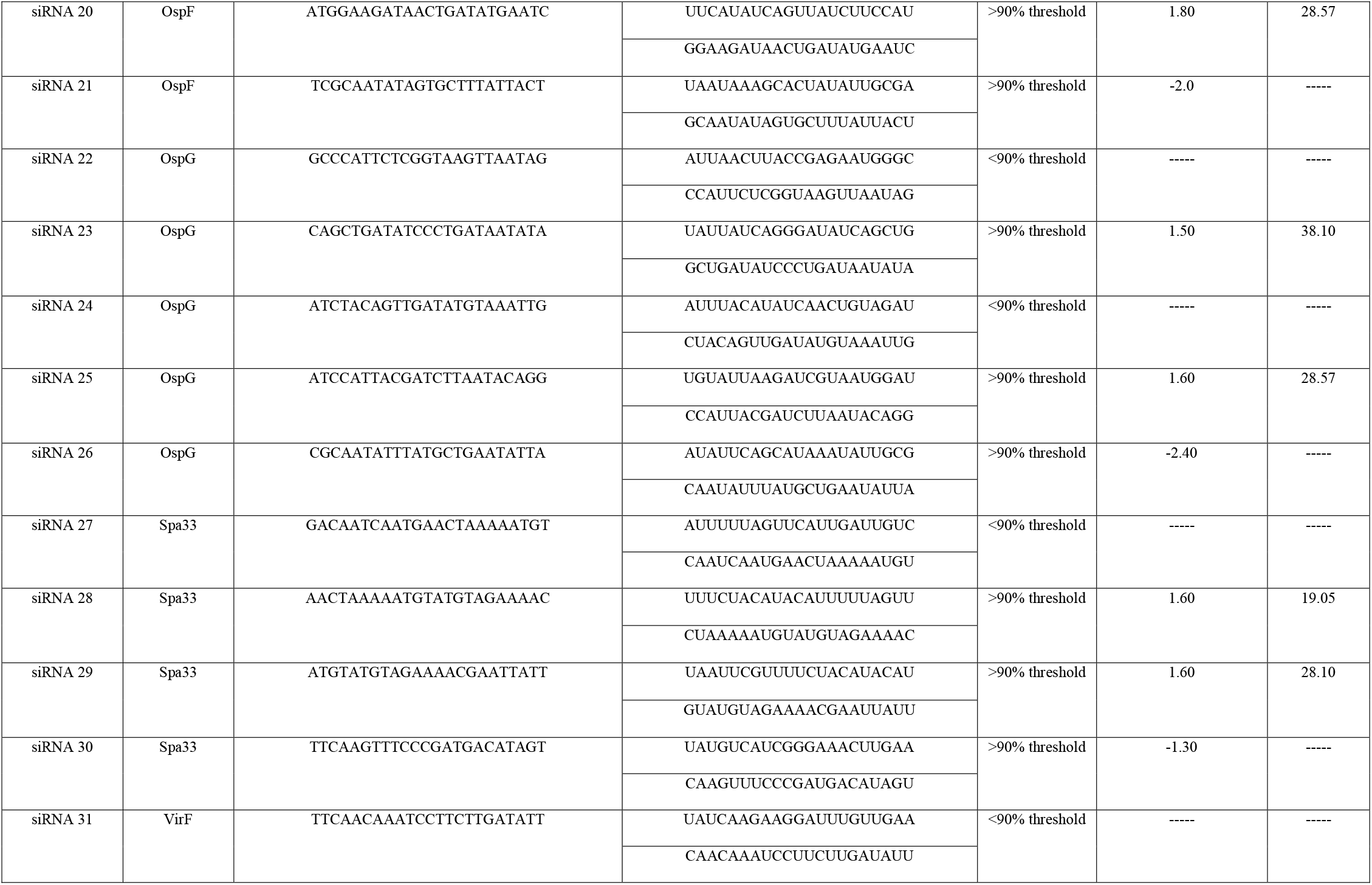
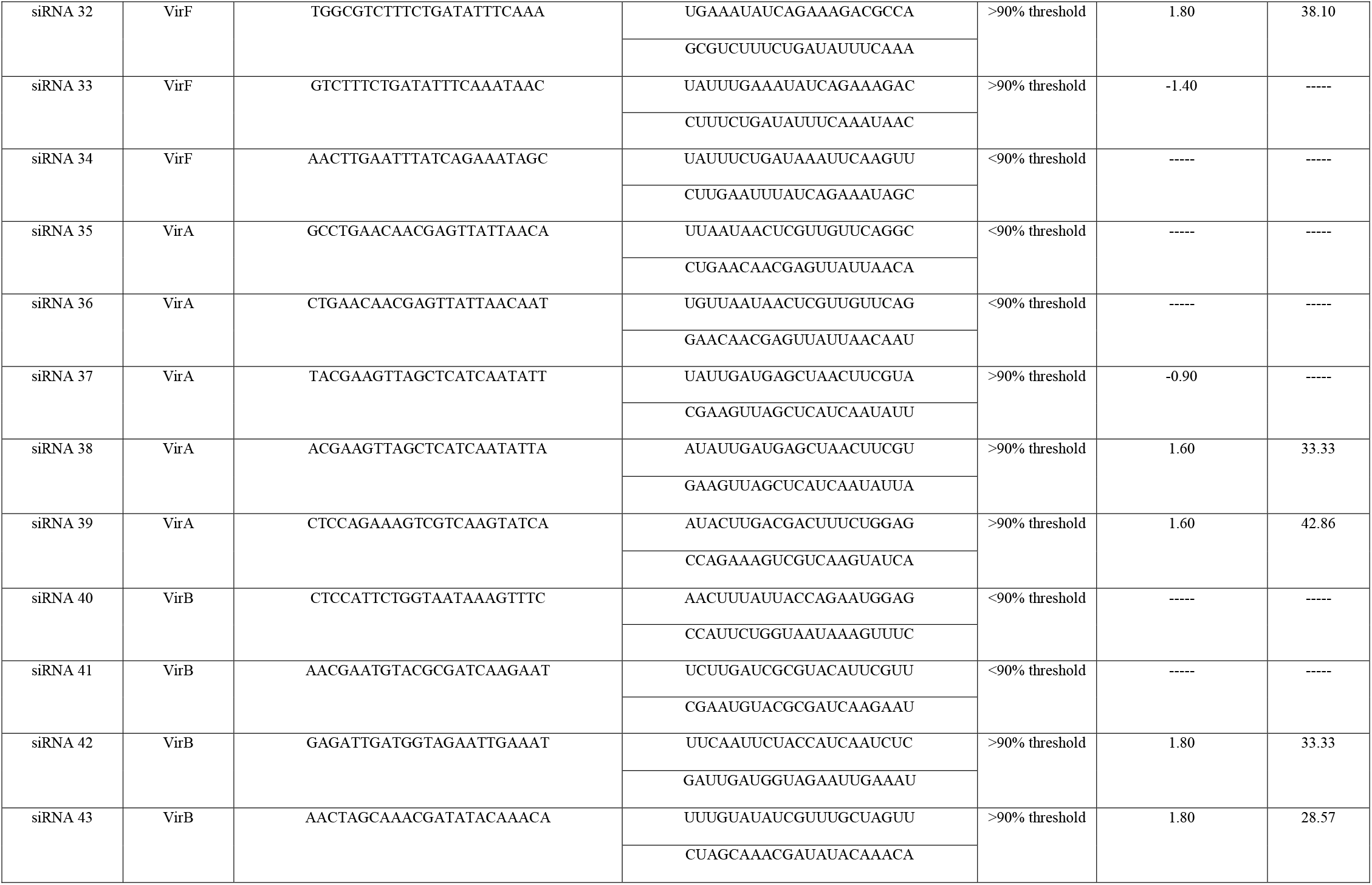
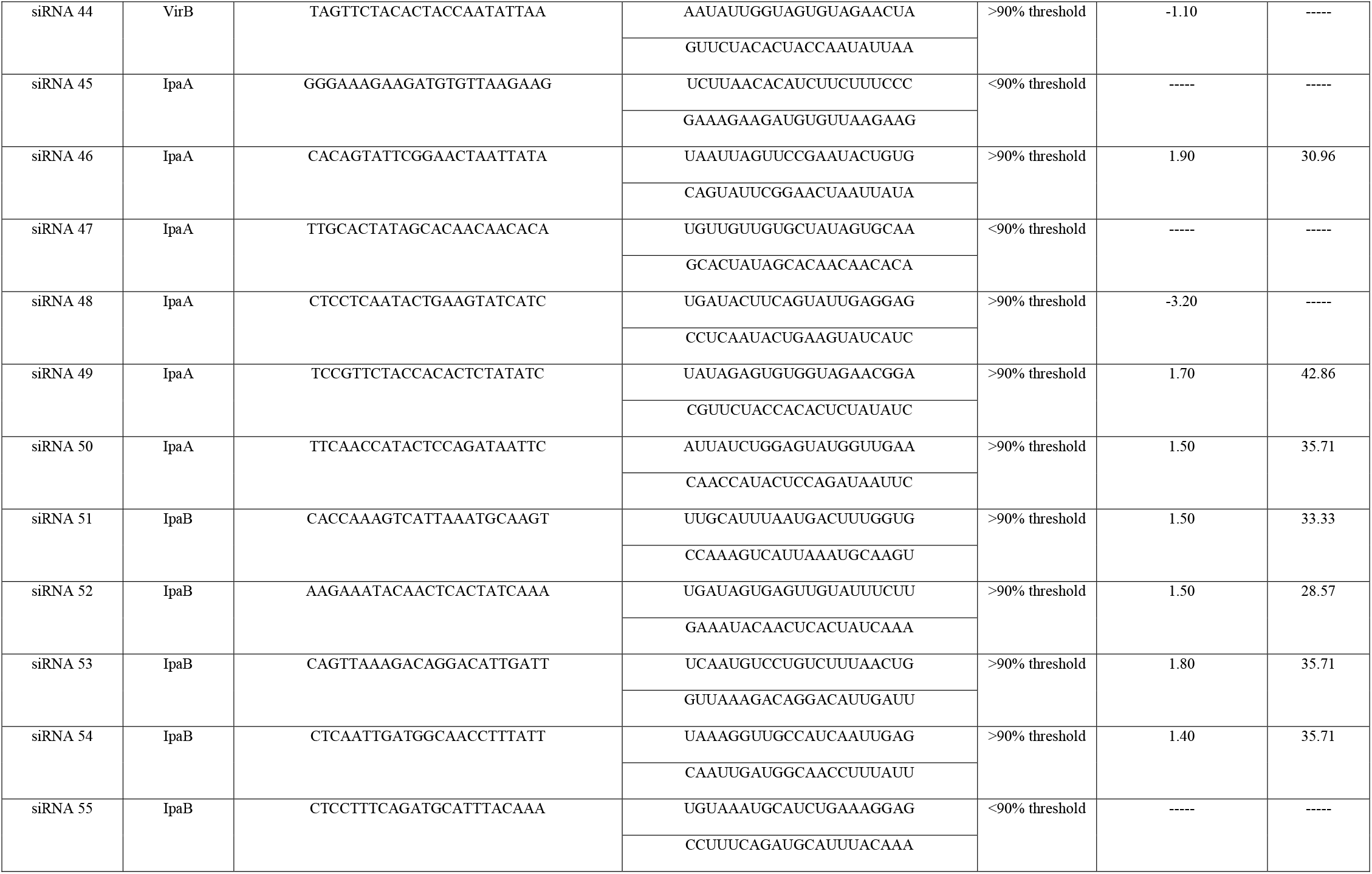
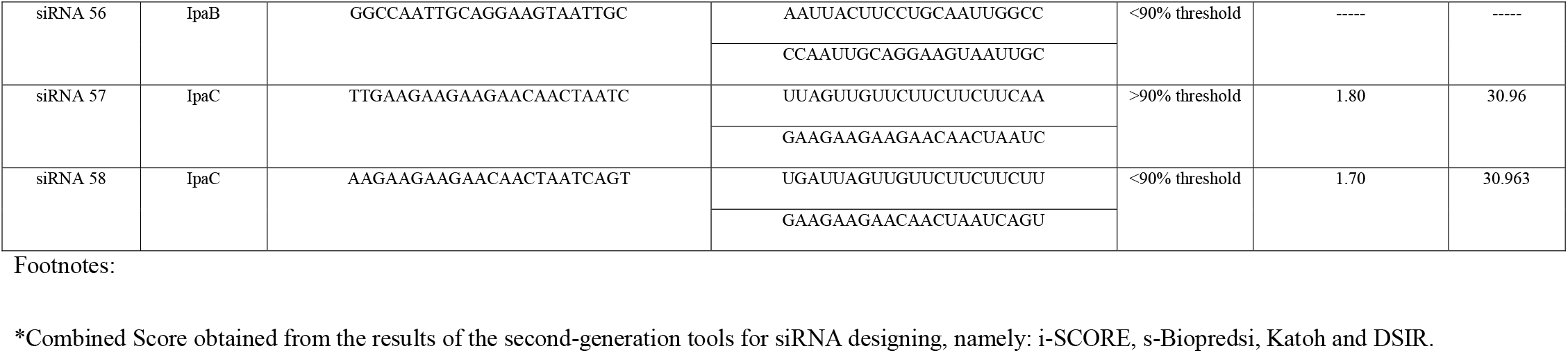
Effective siRNA candidates for each of the conserved virulence associated *Shigella* genes as analyzed by both the first generation and second generation algorithms for designing siRNAs along with their respective free energy of folding and GC content.

### Free energy of self-folding and evaluation of GC content

Functional of siRNA largely depends on molecular structure. For this, extensive efforts have been put on predicting the secondary structure of the of the RNA molecules (49). Benchmark of molecular structural accuracy of the siRNA was set as the “Minimum Free Energy (MFE)” (50). The minimization of free energy is an established phenomenon in computational structural biology on the basis that at an state of equilibrium, the molecule folds into the state of least energy (51).The minimum free energy of folding was calculated by using the RNAstructure web server to check the stability of the predicted siRNA guide strands. From the above mentioned analysis, 30 siRNA candidates were found to yield a positive free energy of folding and thus were considered for further analysis (Table 1).

The GC content of siRNA is a major determinant of the stability of the secondary structure of the siRNA, whereby a GC content ranging between 30-60% is considered sufficient for the execution of its action (52). In our study, out of the 30 siRNA candidates that exhibited a positive free energy of folding in addition to complying with all the primary and secondary algorithms, only 19 candidates were found to have a GC content in between 30-60% and thus were selected for subsequent analysis (Table 1).

### Evaluation of binding energy of siRNA and target and visualization of secondary structures of siRNA-target duplex

Precise prediction of binding energy between siRNA and target is integral for proper understanding of the binding of siRNA to the target and for the subsequent assessment of functional efficiency of the siRNA (53) -(54)-(55). RNA structure, an online web server was used for the estimation of the hybridization energy between the siRNA-target duplex. The thermodynamics of the siRNA-target interaction and the details of the calculation of this binding energy between siRNA and target have been published elsewhere (56)-(57). Table 2 shows the binding energies of the siRNA with the target sequence.

**Table 02:**
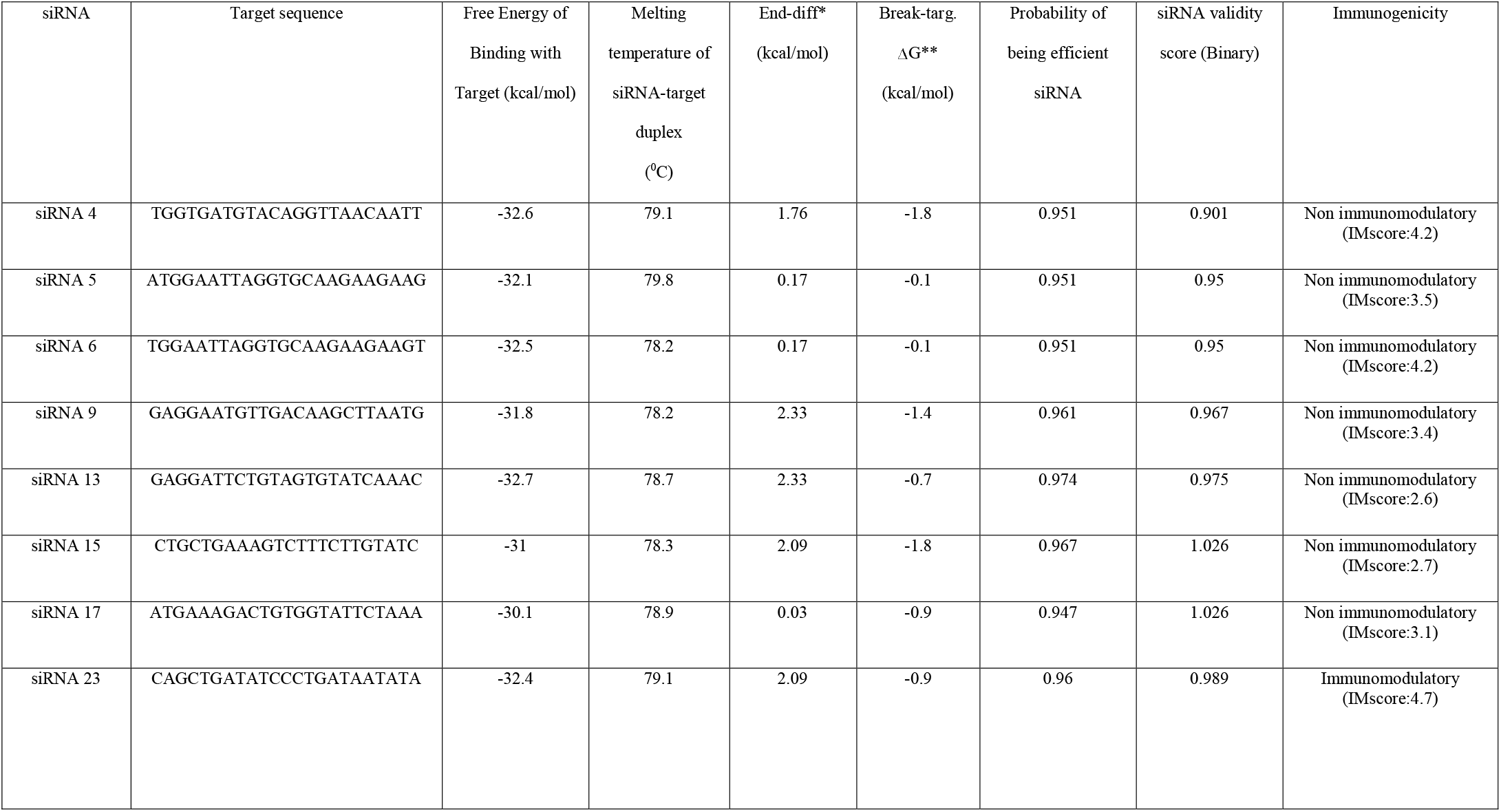
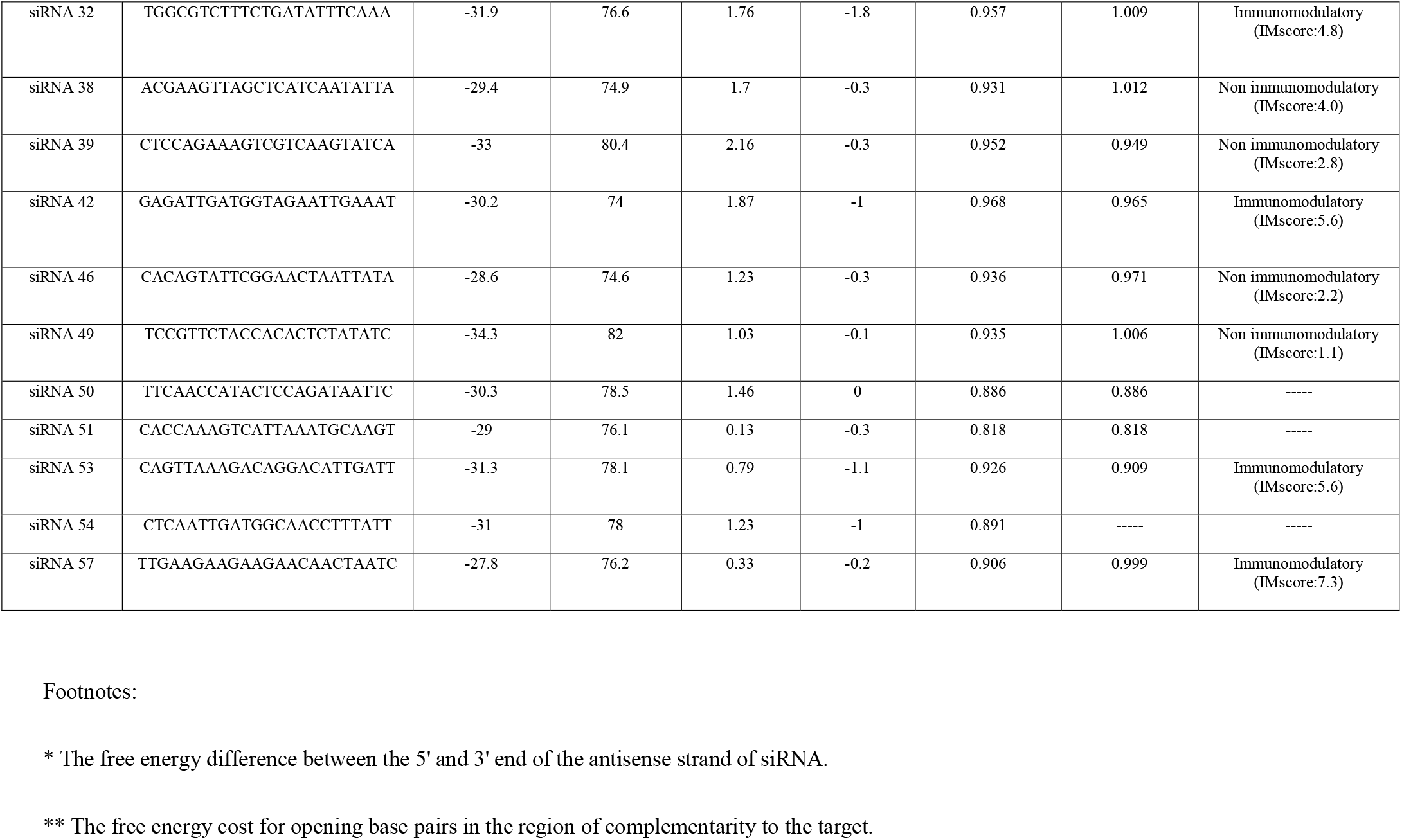
Designed siRNA molecules with their respective free energy of binding of siRNA with target, melting temperature of siRNA-target duplex, target accessibility, functional efficiency, and immunogenicity designed siRNA.

Secondary structures of siRNA with their respective target provide a very efficient computational estimation for both the structure and for the thermodynamics of RNA-RNA interaction (53). RNAstructure program predicts the most stable secondary structures of target-siRNA duplex and minimizes the folding energy. 37 °C is the temperature that was chosen to predict the folded structure. The secondary duplex of candidate siRNA molecules and their corresponding targets are elucidated in **Figure 2**.

### Determination of heat stability of siRNA-target duplex

Heat stability analysis of the siRNA-target duplex, a key factor in the assessment of the stability of secondary structure and functional efficiency of siRNA (58), was conducted by using the OligoWalk server. The results of the heat stability analysis of siRNA-target duplex showed that all the potential 19 siRNA candidates that had fulfilled all the criteria for effective siRNA assessed so far, (as shown in Table 1) had a melting temperature of greater than −70°C (Table 2). Henceforth, the melting temperatures of each of the siRNA-target duplex structure were found to be substantially greater than the physiological temperature of 37.4°C indicating towards the maintenance of integrity of secondary structure of siRNA in the host physiological system.

### Prediction of functional efficiency and target accessibility of the siRNA candidates

Table 2 delineates the summary of the results of the analysis of the functional efficiency and target accessibility as determined by the OligoWalk web server, of each of the siRNA candidates that had fulfilled the threshold for all the criteria set in Table 1. In Table 2, the “End-diff” score for each of the 19 potential siRNA candidates indicates free energy difference between the 5’ and 3’ end of the antisense strand of siRNA (59); whereby the siRNA candidates that have showing positive “End-diff” scores are ranked to have a high functional efficiency. In our study, all the siRNA candidates demonstrated a positive “End-diff” score, thus indicating towards a high functional efficiency of the candidate siRNAs. Moreover, siRNA 9, siRNA 13, siRNA 39, siRNA 15 and siRNA 23 were found to exhibit more positive “End-diff” scores compared to the rest of the siRNA candidates and thus can be classified as having the highest functional efficiency.

Consequently, target accessibility was determined by the “Break targ. ΔG” score, which is free energy account for the opening of base pairs in the region of complementarity to the target (58). Candidate siRNAs demonstrating a less negative “Break targ. ΔG” score are classified as having greater target accessibility (59)-(58). In our study, siRNA 5, siRNA 6, siRNA 49 and siRNA 50 showed the least negative “Break targ. ΔG” score and thus can be classified as having the greatest target accessibility.

Another score delineated in Table 2 as “Probability of being efficient siRNA”, manifests both the functional efficiency and target accessibility of the designed siRNA candidates (58)-(59). In our analysis, siRNA 50, siRNA 51 and siRNA 54 demonstrated a “Probability of being efficient siRNA” score of less than 0.9 and were not considered for further analysis. The rest of the siRNA candidates yield a “Probability of being efficient siRNA” score of greater than 0.9 and thus can be predicted to have high functional efficiency and substantial target accessibility.

### Validation of the functional efficiency of the siRNA candidates

Among the 16 predicted siRNA candidates that had fulfilled the threshold for all the criteria set so far, 11 siRNA candidates (siRNA 4, siRNA 5, siRNA 6, siRNA 9, siRNA 13, siRNA 23, siRNA 39, siRNA 42, siRNA 46, siRNA 53 and siRNA 57) showed a siRNA validity score between 0.8-1.0, following the binary pattern prediction approach indicating high functional efficiency (60). On the other hand, five siRNA candidates (siRNA 15, siRNA 17, siRNA 32, siRNA 38 and siRNA 49) showed a binary validation score of greater than 1.0, which manifests towards the highest functional efficiency (60).

### Prediction of immunotoxicity of the siRNA candidates

We evaluated the immunotoxicity of the 16 siRNA candidates for which functional efficiency was validated in the previous step. 12 out of the 16 siRNA candidates were found to be non-immunogenic on the basis of the default threshold IMscore set for assessing potential immunogenicity of query siRNA sequences and were considered for subsequent analysis. siRNA 23, siRNA 32, siRNA 42 and siRNA 53 were found to be immunogenic (IMscore greater than the threshold of 4.5) and were discarded.

### Designing of tertiary structure of the siRNA candidates and validation

Respective tertiary/3D structures for each of the 12 non-immunogenic siRNA candidates was generated using the web-based RNA modeling software, RNAComposer (61) and the individual 3D structures were saved in pdb format for downstream application. Subsequently, MOLProbity web server was used to validate the individual tertiary structures of the non-immunogenic siRNA candidates and the 3D structures are then filtered for clash score/the number of serious steric overlaps (>0.4 Å) per 1000 atoms (62). Validation scores for the predicted 3D structures of the siRNA candidates have been shown in **Table 3**.

**Table 03:** Validation scores for the predicted 3D structures of the 12 non-immunogenic siRNA candidates.

| siRNA | Nucleic acid Geometry |  |  |  | All Atom Contacts<br>[Clashscore,<br>Percentile] ** |
| --- | --- | --- | --- | --- | --- |
|  | Probability of wrong sugar<br>puckers*<br>(%) | Bad backbone<br>confirmations*<br>(%) | Bad bonds*<br>(%) | Bad angles*<br>(%) |  |
| siRNA 4 | 0.00 | 0.00 | 0.00 | 0.00 | 36.81, 9<sup>th</sup> |
| siRNA 5 | 0.00 | 0.00 | 0.00 | 0.00 | 30.67, 15<sup>th</sup> |
| siRNA 6 | 0.00 | 0.00 | 0.00 | 0.00 | 36.81, 9<sup>th</sup> |
| siRNA 9 | 0.00 | 0.00 | 0.00 | 0.00 | 19.76, 33<sup>rd</sup> |
| siRNA 13 | 0.00 | 0.00 | 0.00 | 0.00 | 24.1, 23<sup>rd</sup> |
| siRNA 15 | 0.00 | 4.76 | 0.00 | 0.00 | 19.23, 35<sup>th</sup> |
| siRNA 17 | 0.00 | 0.00 | 0.00 | 0.00 | 19.61, 34<sup>th</sup> |
| siRNA 38 | 4.76 | 57.14 | 0.00 | 0.00 | 22.52, 26<sup>th</sup> |
| siRNA 39 | 0.00 | 0.00 | 0.00 | 0.00 | 22.46, 26<sup>th</sup> |
| siRNA 46 | 0.00 | 0.00 | 0.00 | 0.00 | 29.76, 16<sup>th</sup> |
| siRNA 49 | 0.00 | 28.57 | 0.00 | 0.00 | 19.03, 35<sup>th</sup> |

\* 5% threshold is allowed for acceptance of the criteria for a valid 3D structure
\*\* 100<sup>th</sup> percentile is the best among the structures for a comparable resolution, 0<sup>th</sup> percentile ranks to be the worst. clashscore is the parameter that represents a comparable resolution.

In our study, a total of three siRNA candidates, namely: siRNA 9, siRNA 15 and siRNA 17 showed an acceptable tertiary structure with all the scores of the analyzed criteria for nucleic acid geometry such as: probability of wrong sugar puckers, bad backbone confirmations, bad bonds and bad angles being below the 5% threshold of acceptance for tertiary structures. **Figure 3** illustrates the tertiary structures of siRNA 9, siRNA 15 and siRNA 17 as visualized using the PyMol Molecular Graphics System (v1.8.4).

## Discussion

Our computational study is concurrent with the current efforts in the development of new therapeutic strategies to combat the persistent conundrum surrounding the emergence of multi-drug resistance among *Shigella* sp. To the best of our knowledge this is the first study that has envisaged proposing a novel treatment option for shigellosis by targeting the virulent proteins which have been reported to be conserved across all known isolates of *Shigella* sp.

In our study, we have used the designing of siRNA candidates that are expected to target the expression of the specific conserved virulence genes of *Shigella* sp through the mediation of a process known as gene silencing (63). Design of siRNA candidates represents a new therapeutic strategy aimed to induce RNA specific inhibition (64), whereby an effective siRNA must fulfill all the threshold criteria set by the first-generation (Ui-Tei, Amarzguioui and Reynolds rules) and second-generation (i-SCORE, s-Biopredsi, DSIR and Katoh rules) algorithms for siRNA designing (65, 66). In this current study, for all the 15 aforementioned conserved virulence genes of *Shigella* sp, we had obtained a total of 58 siRNA candidates that fulfilled the first-generation algorithms, among which only 38 siRNA (Table 1) candidates satisfied all the second-generational algorithms. Consequently, off-target silencing of siRNAs can potentially lead to undesired toxicities (67) and none of the siRNA candidates that satisfied both the first and second generation algorithms were found to exhibit such off-target activity within the human genome.

The folded secondary structure of siRNA is integral for assessment of functional efficiency of siRNA (68) and for the evaluation of structural stability and accuracy of RNA, minimum free energy (MFE) is considered as a standard parameter (50). In our study, out of all the siRNA candidates satisfying both first- and second-generation algorithms, 30 siRNA candidates (Table 1) showed a positive MFE value, indicating thermodynamic infeasibility of self-folding. Subsequently, only 19 of these 30 siRNA candidates with a positive MFE value were found to have GC content within the range of 30-60%. GC content between 30% and 60% is recommended for siRNA sequence, since there is a considerable reciprocal correlation between GC-content and RNAi activity (52), whereby a GC content of siRNA within the aforementioned range exhibits stronger inhibitory effect (69).

Prediction of secondary structure between siRNA and target mRNA acts as an integral cue to the selection of specific siRNA target site (69). Random folding of siRNA may lead to the inhibition of its RNAi activity with an inappropriate secondary structure of siRNA-duplex can result in impediment to the RISC (RNA-induced silencing complex) formation (70). Henceforth, evaluation of the thermodynamic outcome for the interaction between the siRNA candidate and the target mRNA involves the assessment of the sum of the energy required to unravel the binding site and the energy gained from the resultant hybridization process (56). In our study, all the 19 siRNAs that were analyzed for feasibility in binding to target mRNA exhibited highly negative ΔG values for binding to target sequences, thus indicating thermodynamically feasible target binding.

The designing of therapeutic siRNA candidates involves the analysis of heat stability of the siRNA for the evaluation of its in vivo stability and functional efficiency (71). In our study, all the 19 siRNA candidates analyzed for heat stability were found to exhibit melting temperatures considerably greater than the physiological temperature, thus indicating structural integrity in the host system. Consequently, assessment of target accessibility of the siRNA candidates indicates the efficiency of efficiency of RISC mediated endonucleolytic cleavage, which is the final step in the biological mechanism of gene silencing by siRNA (72). siRNA 5, siRNA 49 and siRNA 50 exhibited the best scores for assessment of target accessibility; i.e-least negative “Break targ. ΔG” values.

The immune system is armed with the required machinery to recognize foreign RNA sequences, resulting in the mediation of activation of pattern recognition regions (PRR) for the clearance of the exogenous components (73). Thereby, the immunogenicity of an RNA sequence in the case of siRNA-based therapy may lead to immunotoxicity (74). Our assessment of immunogenicity of the candidate siRNAs showed that a total of 12 siRNAs were non-immunogenic (Table 2) and thus were considered for tertiary structure validation. Owing to the small number of siRNA molecules being evaluated by X-ray crystallography, RMN spectroscopy, and cryoelectronic microscopy (cryo-EM), tertiary structure prediction of siRNA is integral in understanding the respective structure-function relationship (75). Among the 12 siRNA candidates that were analyzed for tertiary structure validation, only three siRNA candidates (siRNA 9, siRNA 15 and siRNA 17) satisfied all threshold scores for nucleic acid chemistry parameters such as: RNA sugar puckers, RNA backbone conformations and bond angles (Table 3).

Henceforth, the siRNA candidates that were found to have an acceptable tertiary structure are intended to halt the translation of distinct conserved virulence genes of *Shigella* sp. siRNA 9 is intended to bind to the mRNA of Ipg D gene, in the process halting the expression of the IpD protein that is involved with the entry of the bacteria through ruffling of the host membrane (34). Thus, siRNA 9 is expected to protect the host system through the maintenance of host membrane integrity. siRNA 15 and siRNA 17 are intended to target the expression of the OspB gene, thereby halting the process of host inflammatory response and resulting tissue damage (34).

## Limitations of the study

Despite the long-term prospects of the findings intended for the development of a novel therapeutic strategy against shigellosis, our study has several limitations. First of all, our analysis is solely based on the sequences of the isolates reported in the NCBI database and may not necessarily be applicable for isolates with novel mutations in these conserved virulence genes that may be reported in the future. Moreover, the implementation of siRNA in regular therapeutic applications is a distant prospect and requires extensive wet lab validation regarding the mode of delivery of the siRNA into the host system as well as modifications in the siRNA sequences may be required for enhanced efficacy in the host system.

## Conclusion

In this study, we have proposed three distinct siRNA candidates that were found to have substantial functional efficacy, non-immunogenic and have an acceptable tertiary structure that is integral for the effectiveness of siRNA in the host system. Our study provides insights into the development of a new form of therapeutics against shigellosis in the form of siRNA. Such future therapeutic strategies may have promising implications in combating the rapid development of anti-microbial resistance among *Shigella* sp and the emergence of multi-drug resistant isolates.

## Methods and Materials

An overview of the methodology followed in our study is illustrated by **Figure 1**.

**Figure 1.**
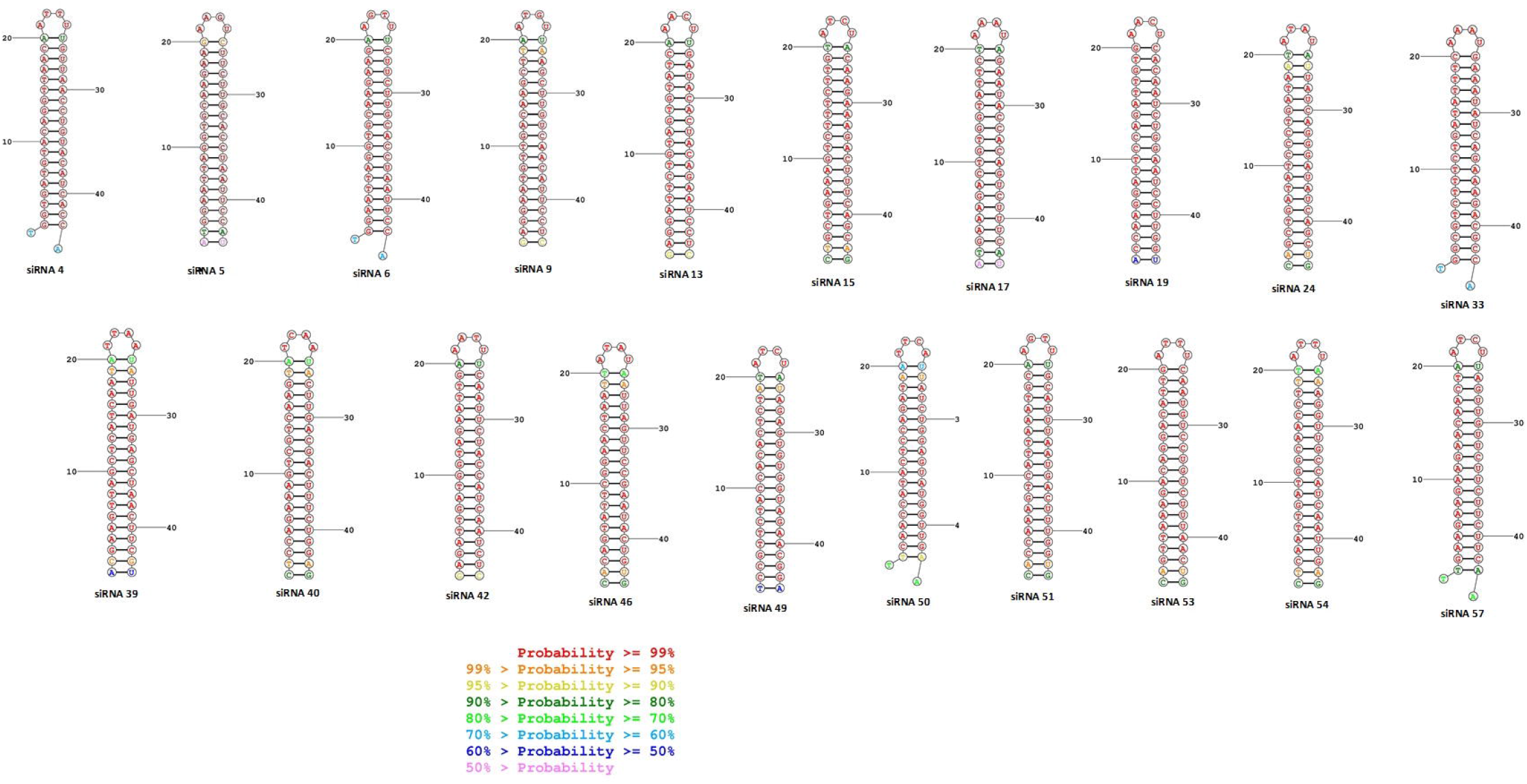
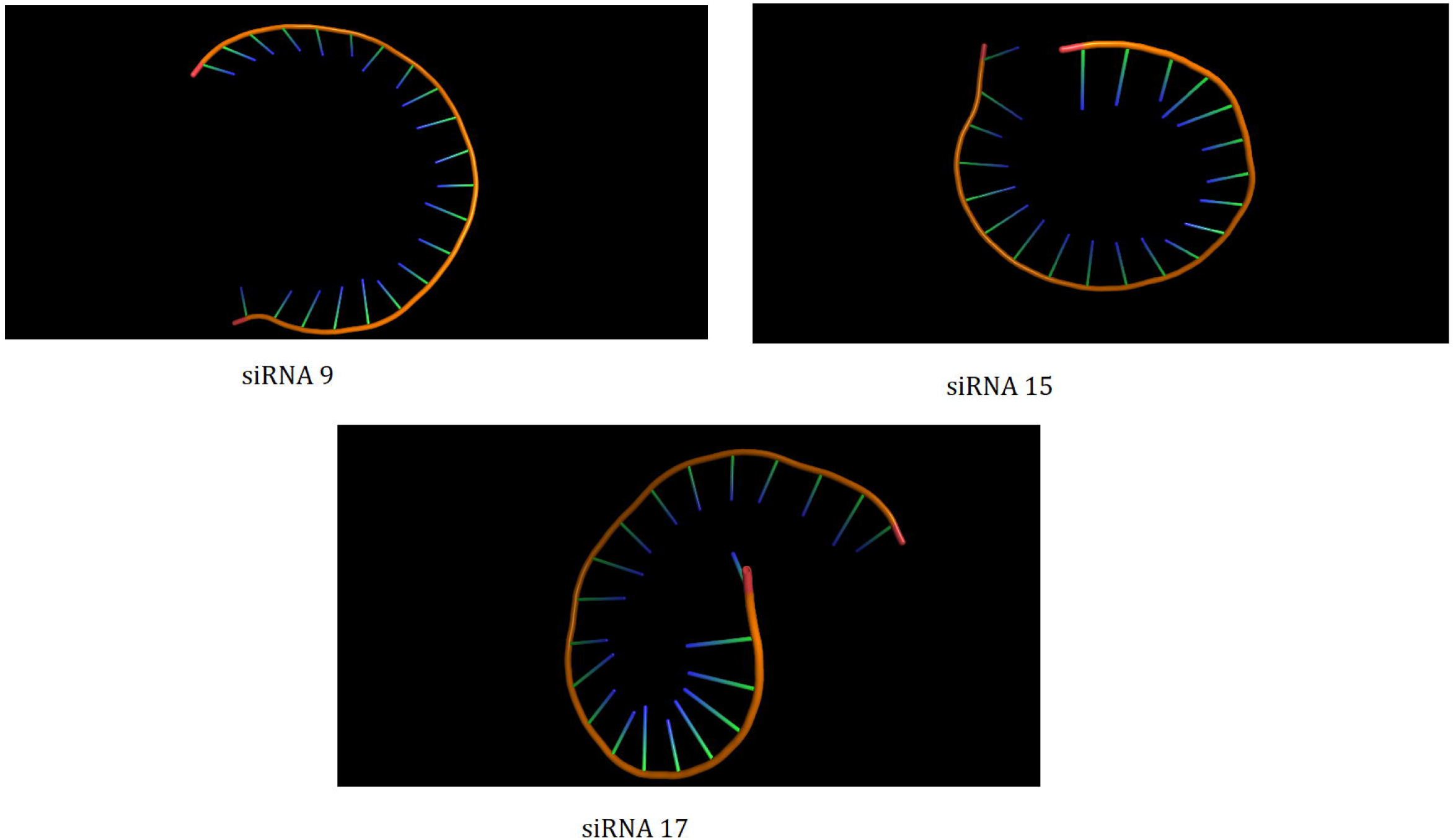
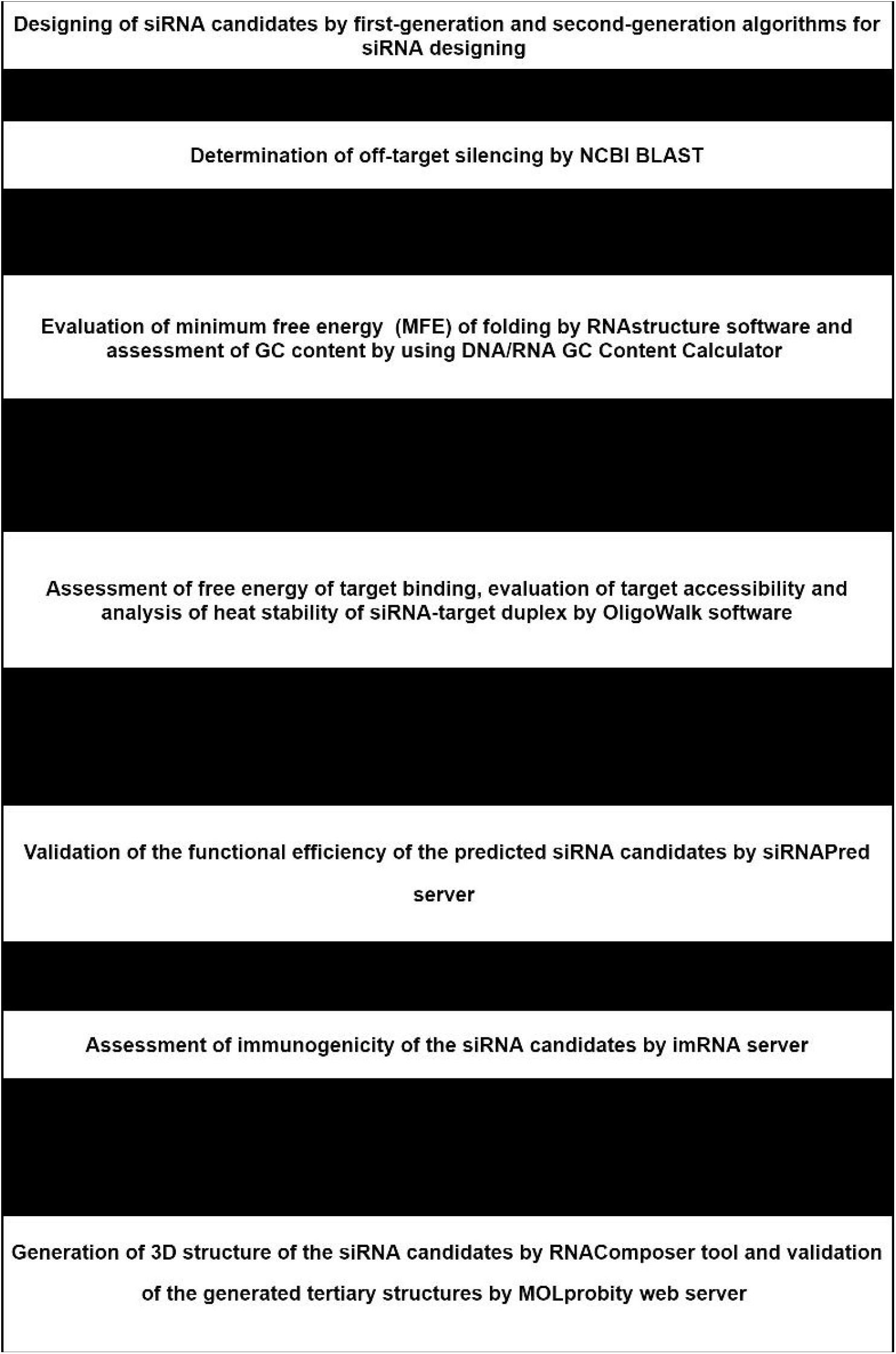
Overview of the workflow of the methodology followed in the study.

### Sequence retrieval and multiple sequence alignment for determination of conserved regions

Complete gene sequences for 15 different conserved virulent proteins of multiple isolates of *Shigella* sp. were obtained from NCBI Nucleotide (https://www.ncbi.nlm.nih.gov/nucleotide). These conserved virulent proteins of *Shigella* sp. included: IcsA (from 40 isolates), IpaA (from 64 isolates), IpaB (from 55 isolates), IpaC (from 41 isolates), IpaJ (from 57 isolates), IpgB (from 46 isolates), IpgD (from 42 isolates), MxiC (from 62 isolates), OspB (from 68 isolates), OspF (from 28 isolates), OspG (57 isolates), Spa33 (from 21 isolates), VirA (from 50 isolates), VirB (from 61 isolates) and VirF (68 isolates). Identification of conserved regions from these 15 different virulent proteins of *Shigella* sp was done by multiple sequence alignment using Clustal Omega (https://www.ebi.ac.uk/Tools/msa/clustalo/).

### Recognition of target sequence and des igning of potential siRNA candidates

For the purpose of identification of target sequence and siRNA designing, siDirect 2.0 (http://sidirect2.rnai.jp/) an efficient and target-specific siRNA designing tool was used (47). This tool employs a combination of first-generation algorithms for siRNA designing, namely: Ui-Tei, Amarzguioui, Reynolds rules (URA rules) along with a melting temperature of less than 21.5 °C as the absolute parameters for prediction of potential siRNA duplex formation (76). Moreover, a multitude of other components that were also taken into consideration on the concept of the URA rules as shown in Supplementary Table 1 have been used in a number of previously published literatures (42).

The i-SCORE Designer, an online software (48) was used for the validation of the potential siRNA candidates obtained from siDirect 2.0. This web-based software utilizes multiple second generation algorithms in addition to the first-generation algorithms for siRNA designing (U, R, A rules), among which i-SCORE, s-Biopredsi, Katoh and DSIR rules are the prominent second-generation algorithms (77)-(78)-(79). Only those siRNA candidates that were found to fulfill the threshold set by the criteria for the second-generation algorithms were subsequently selected for further downstream validation.

### Determination of off-target similarity

Off-target sequence similarity for the guide strands of the siRNA candidates was checked using the BLAST tool (http://www.ncbi.nlm.nih.gov/blast) against the entire GenBank database with default threshold value of 10 and BLOSUM 62 as parameter.

### Prediction of free energy of folding and calculation of GC content

The free energy of folding of the guide strand of the siRNA candidates was assessed using RNAstructure (https://rna.urmc.rochester.edu/RNAstructureWeb/), a web-based tool for prediction of secondary structure of RNA (80). Only those siRNA candidates that had exhibited a positive free energy of folding (positive ΔG) were used for the subsequent process of GC content calculation by using DNA/RNA GC Content Calculator (http://www.endmemo.com/bio/gc.php) and siRNA candidates with a GC content of 30-60% were subsequently selected for downstream validation.

### Evaluation of the thermodynamics secondary structure formed between siRNA and target and visualization of siRNA-target binding

Investigation of the thermodynamics for the secondary structure formed between the guide strand and subsequent validation of the siRNA-target duplex was done using the web-based tool for RNA secondary structure prediction, RNAStructure (https://rna.urmc.rochester.edu/RNAstructureWeb/). This tool determines the hybridization energy and base pairing from two RNA sequences by following the functional algorithm of McCaskill’s partition to compute probabilities of base pairing, realistic communication energies and equilibrium concentrations of duplex structures (80)-(81).

### Determination of heat stability and prediction of functional efficiency and target accessibility of the potential siRNA candidates

For the determination of heat stability of the guide strand as well as for the assessment of functional efficiency and target accessibility of the predicted siRNA candidates, we used the features of OligoWalk (http://rna.urmc.rochester.edu/cgi-bin/server_exe/oligowalk/oligowalk_form.cgi), a web-based server for calculating thermodynamic features of sense-antisense hybridization (82). This tool operates through a designated query involving the sequence for the guide strand of the siRNA candidate and the subsequent results obtained are expressed as: “End-diff (free energy difference between the 5’ and 3’ end of the antisense strand of siRNA)”, which indicates denotes the functional efficiency of the siRNA candidate; “Break-targ. ΔG (free energy cost for opening base pairs in the region of complementarity to the target)”, which signifies the target accessibility of the designed siRNA and “Probability score of being efficient siRNA”, which is calculated on the basis of both target accessibility and functional efficiency of the designed siRNA (82)-(59).

### Validation of the functional efficiency of the siRNA candidates

siRNAPred server (http://crdd.osdd.net/raghava/sirnapred/index.html) was used for the validation of the functional efficiency of the predicted siRNA candidates against the Main21 dataset using support vector machine algorithm and the binary pattern prediction approach (60). Validation scores from the server greater than 1 predicts very high efficiency, scores ranging from 0.8-0.9 predict high efficiency and scores ranging from 0.7-0.8 predicts moderate efficiency (60).

### Prediction of immunotoxicity of the predicted siRNA candidates

For the purpose of prediction of immunotoxicity of the designed siRNA candidates, imRNA (https://webs.iiitd.edu.in/raghava/imrna/sirna.php), a web-based server consisting of various components integral for the designing of RNA-based therapeutics was employed (83). IMscore of 4.5 set as default was used for the prediction of immunogenicity of potential siRNA candidates.

### Designing of tertiary (3D) structure of the siRNA candidates and validation

Designed siRNA candidates that had surpassed the immunogenicity filter set in the previous step were subject to tertiary structure prediction using RNAComposer (http://rnacomposer.cs.put.poznan.pl/), an automated tool that predicts the tertiary structure from a linear siRNA sequence (84)-(85). Subsequently, the tertiary structures of these siRNA candidates were validated using the MOLprobity web server (http://molprobity.biochem.duke.edu/) (62), which uses the pdb file of the tertiary structure of the siRNA as input to predict the validity of the tertiary structure on the basis of all-atom contacts and geometry, RNA sugar puckers, RNA backbone conformations, hydrogen bonds and Van der Waals forces (62)-(86). The tertiary structures of the candidate siRNAs with an acceptable tertiary structure were visualized using the PyMOL Molecular Graphics System (v1.8.4).

## Supplementary Information

**Supplementary File 1. List of conserved sequences from the 15 virulent genes of *Shigella* sp.**

**Supplementary Table 1. Additional rules applied along with the URA rules for siRNA designing.**

## Acknowledgments

The authors thank Mohammad Umer Sharif Shohan, Lecturer, Department of Biochemistry and Molecular Biology, University of Dhaka for his insightful suggestions during the work. ICDDR,B is grateful to the Government of Bangladesh, Canada, Sweden, and the UK for providing unrestricted core support. The authors gratefully acknowledge these donors for their support and commitment to the endeavors of ICDDR,B.

## Author Contributions

P.P, N.B and Z.N conceived the study; P.P and F.T.C designed the study; B.S, S.N.M, M.U.S.S and U.F.C were involved in the acquisition of the data; P.P, M.K, T.J.S and S.N were involved in the data acquisition. P.P, T.J.S, Z.N and TA were involved in drafting of the manuscript. All authors read and approved of the manuscript.

## Institutional Review Board Statement

Not applicable.

## Informed Consent Statement

Not applicable.

## Data Availability Statement

All relevant data are contained within this article.

